# Microtubule assembly by soluble tau impairs vesicle endocytosis and excitatory neurotransmission via dynamin sequestration in Alzheimer’s disease synapse model

**DOI:** 10.1101/2021.09.13.460074

**Authors:** Tetsuya Hori, Kohgaku Eguchi, Han-Ying Wang, Tomohiro Miyasaka, Laurent Guillaud, Zacharie Taoufiq, Hiroshi Yamada, Kohji Takei, Tomoyuki Takahashi

## Abstract

Elevation of soluble wild-type (WT) tau occurs in synaptic compartments in Alzheimer’s disease. We addressed whether tau elevation affects synaptic transmission at the calyx of Held in brainstem slices. Whole-cell loading of WT human tau (h-tau) in presynaptic terminals at 10-20 µM caused microtubule (MT) assembly and activity-dependent rundown of excitatory neurotransmission. Capacitance measurements revealed that the primary target of WT h-tau is vesicle endocytosis. Blocking MT assembly using nocodazole prevented tau-induced impairments of endocytosis and neurotransmission. Immunofluorescence imaging analyses revealed that MT assembly by WT h-tau loading was associated with an increased bound fraction of the endocytic protein dynamin. A synthetic dodecapeptide corresponding to dynamin-1-pleckstrin-homology domain inhibited MT-dynamin interaction and rescued tau-induced impairments of endocytosis and neurotransmission. We conclude that elevation of presynaptic WT tau induces *de novo* assembly of MTs, thereby sequestering free dynamins. As a result, endocytosis and subsequent vesicle replenishment are impaired, causing activity-dependent rundown of neurotransmission.

**Significance Statement:** Wild-type human recombinant tau loaded in rodent presynaptic terminals inhibited vesicle endocytosis, thereby causing activity-dependent rundown of excitatory transmission. This endocytic block is caused by a sequestration of dynamin by excess microtubules newly assembled by tau and can be rescued by a peptide inhibiting the microtubules-dynamin interaction, or by the microtubule disassembler nocodazole. Thus, synaptic dysfunction can be induced by pathological increase of endogenous soluble tau in Alzheimer disease slice model.

## Introduction

The microtubule (MT) binding protein tau assembles and stabilizes MTs (1, 2) mainly in axonal compartments (3, 4). Phosphorylation of tau proteins reduces their binding affinity (5, 6), thereby shifting the equilibrium from MT-bound form to soluble free form (7). Soluble tau proteins also exist in dynamic equilibrium between phosphorylated and dephosphorylated forms (8) as well as between soluble and aggregated forms. When the cytosolic tau concentration is elevated, monomeric tau undergoes oligomerization and eventually precipitates into neurofibrillary tangles (NFT) (9-11), which is a hallmark of tauopathies, including Alzheimer’s disease (AD), frontotemporal dementia with Parkinsonism-17 (FTDP-17) and progressive supranuclear palsy (PSP) (2, 7, 8). Although the NFT density can correlate with the degree of AD progression (2, 7, 12), soluble tau protein levels are more closely linked to disease progression and cognitive decline (13, 14).

Genetic ablation of tau shows little abnormal phenotype (15-17), presumably due to compensation by other MT-associated proteins (15). Instead, tau ablation can prevent amyloid β-induced impairments of mitochondrial transport (16) or memory defects (18, 19). Thus, loss of tau function due to its dissociation from MTs is unlikely to be an important cause of neuronal dysfunction in AD (8, 12).

In postmortem brains of both AD patients and intact humans, tau is present in synaptosomes (20, 21). In a transgenic mice AD model, soluble tau is accumulated in the hippocampal nerve terminal zone (22, 23). Both *in vivo* and in culture models of tauopathy, tau is released from axon terminals upon KCl stimulation in a Ca^2+^-dependent manner, like neurotransmitters (24, 25). Tau oligomers produced by released tau triggers endogenous tau seeding in neighboring neurons, thereby causing trans-synaptic propagations (22, 26).

FTDP tauopathy model mice that overexpressed with mutant tau are widely used to examine tau toxicities on synaptic plasticity (27-29), memory formation (28, 30) as well as on synaptic vesicle transport (31, 32). In contrast to FTDP, which is a rare familial disease associated with tau mutation, AD is a widespread sporadic disease unassociated with tau mutation, but the expression level of WT tau being crucial. As AD models, the effects of WT tau overexpression have been examined in culture cells (33-36) or in *Drosophila* (37), where impaired axonal transports associated with increased MT density were found. These observations suggest that WT tau can be detrimental when its levels are elevated (35, 36). However, unlike FTDP tau mutant, it is unknown whether elevated soluble WT tau can affect mammalian central synaptic transmission.

We addressed this question using the giant nerve terminal calyx of Held visualized in slices from mice brainstem, where axonal MTs extended into the depth of terminals (38). In this presynaptic terminal, we loaded recombinant WT h-tau from a whole-cell patch pipette at fixed concentrations to model the elevation of WT tau associated with AD and found that WT h-tau newly assembled MTs and strongly impaired synaptic transmission. Capacitance measurements indicated that the primary target of WT h-tau is vesicle endocytosis. Immunocytochemical image analysis after cell permeabilization revealed an increase in the bound fraction of the endocytic GTPase dynamin in WT h-tau-loaded terminals. Since the endocytic key protein dynamin is a MT-binding protein (39), dynamin is likely sequestered by newly assembled MTs. Out of screening, we found that a synthetic dodecapeptide corresponding to amino acids 560-571 of dynamin 1 inhibited MT-dynamin interaction. When we co-loaded this peptide “PHDP5” with WT h-tau, its toxicities on vesicle endocytosis as well as on synaptic transmission were rescued. Thus, we propose a novel synaptic dysfunction mechanism underlying AD, in which WT tau-induced over-assembly of MTs depletes dynamins, thereby impairing vesicle endocytosis and synaptic transmission.

## Results

### Intra-terminal loading of WT h-tau impairs excitatory synaptic transmission

To address whether elevation of soluble h-tau in presynaptic terminals can affect synaptic transmission, we purified WT recombinant h-tau (0N4R) and its deletion mutant (del-MTBD) lacking the MT binding site (244Gln-367Gly) (Figure S1A), obtained using an *E. coli* expression system (40). These recombinant h-tau proteins are highly soluble at room temperature without any sign of granulation (41). In simultaneous pre- and postsynaptic recording at the calyx of Held in mouse brainstem slices, we recorded EPSCs evoked at 1 Hz by presynaptic action potentials (Figure 1). After confirming stable EPSC amplitude for 10 min, we injected a large volume of internal solution containing WT h-tau (20 µM) from an installed fine tube to a presynaptic whole-cell pipette to replace most of pipette solution and allow h-tau to diffuse into a presynaptic terminal (illustration in Figure 1A) (42, 43). After loading h-tau (20 µM), the amplitude of glutamatergic EPSCs gradually declined and reached 23 ± 9 % in 30 min (Figure 1A, p < 0.01, paired t-test, n = 6 synapses in 6 slices). WT h-tau loaded at a lower concentration (10 µM) caused a slower EPSC rundown to 65 ± 5 % in 30 min (p < 0.01, n = 5 synapses in 5 slices). Loading of del-MTBD (20 µM), lacking tubulin polymerization capability (Figure S1B), likewise loaded had no effect on EPSC amplitude (Figure 1A). Since h-tau concentrations in presynaptic terminals are equilibrated with those in a presynaptic whole-cell pipette with a much greater volume than terminals (44), these results suggest that WT h-tau >10 µM can significantly impair excitatory synaptic transmission.

**Figure 1.**
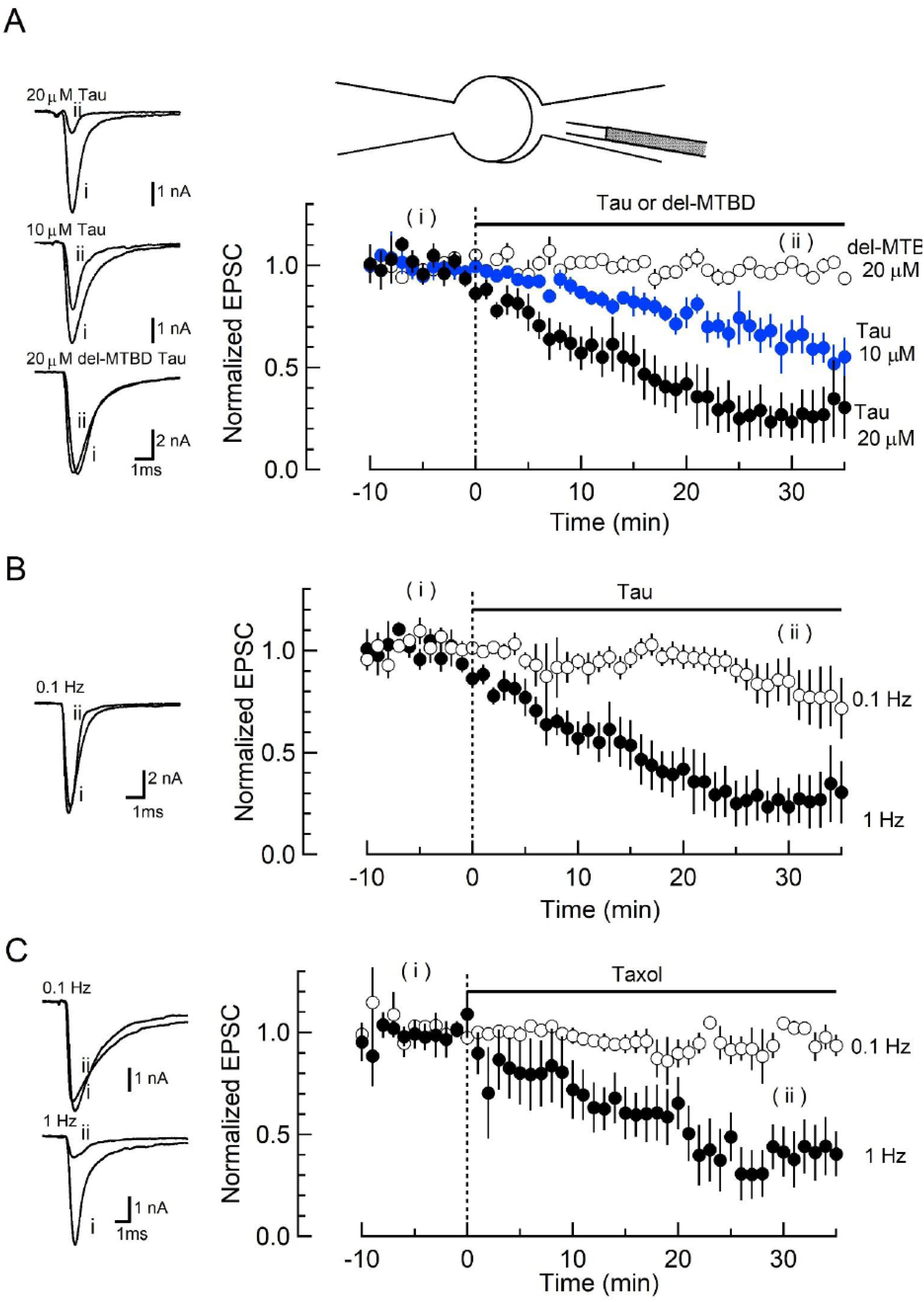
WT h-tau loaded in presynaptic terminals inhibited excitatory synaptic transmission. (A) In simultaneous pre- and postsynaptic whole-cell recordings, intra-terminal infusion of WT h-tau at 10 µM (blue filled circles) or 20 µM (black filled circles), from a tube in a presynaptic patch pipette (top illustration), caused a concentration-dependent rundown of EPSCs evoked by presynaptic action potentials at 1 Hz. In the time plots, EPSC amplitudes averaged from 60 events are sampled for data points and normalized to the mean amplitude of baseline EPSCs before h-tau infusion. Sample records of EPSCs 5 min before (i) and 30 min after (ii) tau infusion are superimposed and shown on the left panels. The EPSC amplitude remaining 30 min after infusion was 23 ± 9 % and 65 ± 5 %, respectively for 10 µM and 20 µM h-tau (means and SEMs, 6 synapses from 6 slices, p < 0.01 in paired t-test between before and after h-tau infusion). Infusion of MT-binding site-deleted h-tau mutant (del-MTBD, 20 µM, Supplemental Figure S1A) had no effect on the EPSC amplitude (open circles, sample EPSC traces shown on the left bottom panel). (B) The amplitude of EPSCs evoked at 0.1 Hz remained unchanged after h-tau infusion (85 % ± 12 %, 5 synapses from 5 slices, p = 0.22 in paired t-test). Sample records of EPSCs before (i) and 30 min after (ii) h-tau infusion at 0.1 Hz are superimposed on the left panel. (C) Taxol (1 µM) caused activity-dependent rundown of EPSC amplitude to 41.4 ± 12 % at 1 Hz (p < 0.01, 5 synapses from 5 slices), but remained unchanged when stimulated at 0.1 Hz (105 ± 3.0 %, open circles, 4 synapses from 4 slices). Sample records of EPSCS at 0.1 Hz and 1 Hz are superimposed on the left panels.

The inhibitory effect of WT h-tau on EPSCs was apparently frequency-dependent. When evoked at 0.1 Hz, WT h-tau (20 µM) caused only a minor reduction of EPSC amplitude (85 ± 12 %, 30 minutes after loading, p = 0.21, n = 5; Figure 1B). Since taxol shares a common binding site of MTs with tau (45) and assembles tubulins into MTs (Figure S1), we tested the effect of taxol (1 µM) on EPSCs (Figure 1C). Like h-tau, taxol caused a significant rundown of EPSCs evoked at 1 Hz (to 41 ± 12 at 30 min, n =5, p< 0.05), but not those evoked at 0.1 Hz (104 ± 3.0 %, n = 4, p = 0.60). These results together suggest that MTs newly assembled in presynaptic terminals by WT h-tau or taxol cause activity-dependent rundown of excitatory synaptic transmission.

### WT h-tau primarily inhibits SV endocytosis and secondarily exocytosis

To determine the primary target of h-tau causing synaptic dysfunction, we performed membrane capacitance measurements at the calyx of Held (46-49). Since stray capacitance of perfusion pipettes prevents capacitance measurements, we backfilled h-tau into a conventional patch pipette after preloading normal internal solution only at its tip to secure GΩ seal formation. This caused substantial and variable delays of the intra-terminal diffusion, so no clear effect could be seen more than 10 minutes after whole-cell patch membrane was ruptured. 20 minutes after patch-loading of WT h-tau (20 µM), endocytic capacitance showed a significant slowing (Figure 2), whereas exocytic capacitance magnitude (ΔC_m_) or charge of Ca^2+^ currents (Q_Ca_) induced by a depolarizing pulse was not different from controls without h-tau loading. 30 minutes after loading h-tau, the endocytic capacitance change became further slowed (p < 0.01), and exocytic ΔC_m_ eventually showed a significant reduction (p < 0.05, n = 5) without a change in Q_Ca_. These results suggest that the primary target of h-tau toxicity is synaptic vesicle (SV) endocytosis. Endocytic block inhibits recycling replenishment of SVs via recycling, thereby reducing the exocytic release of neurotransmitter as a second effect.

**Figure 2.**
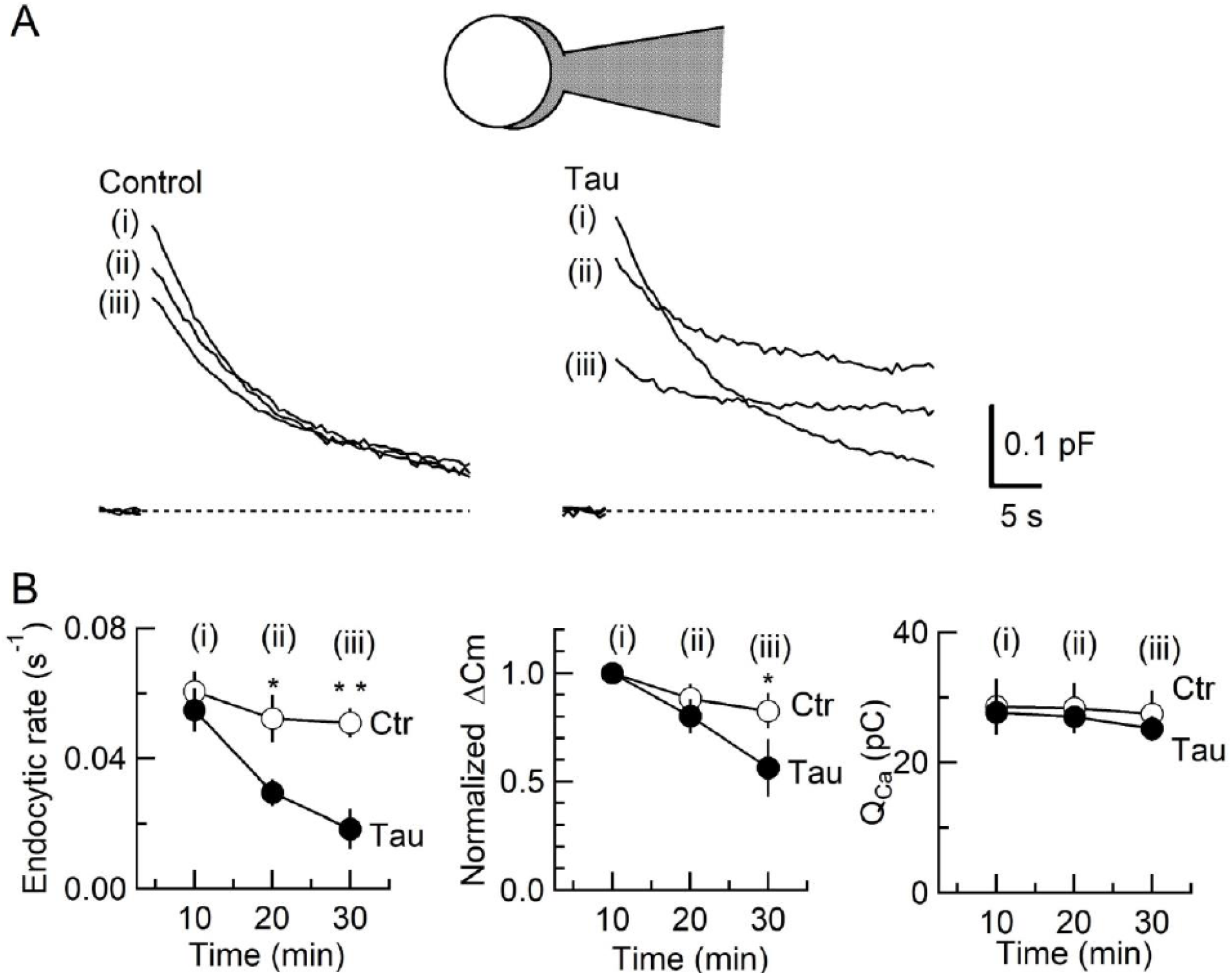
Inhibition of SV endocytosis is the primary effect of WT h-tau loading. (A) Exo-endocytic membrane capacitance changes in presynaptic terminals without (Control) or after direct loading of WT h-tau (20 µM; Tau). WT h-tau was directly loaded by diffusion into a terminal from a whole-cell patch pipette (illustration). Capacitance traces were sampled from (i) 10, (ii) 20 and (iii) 30 min after patch membrane rupture (superimposed). *Left panel*, non-loading control. *Right panel*, WT h-tau-loaded terminal. Capacitance changes were evoked every 2 min by Ca^2+^ currents induced by a 20-ms depolarizing pulse (not shown). (B) Time plots of endocytic rate (left panel), exocytic magnitude (middle panel) and presynaptic Ca^2+^ current charge (right panel). Data points represent averaged values from 5 events from 4 min before and 4 min after the time points. In calyceal terminals, 20 min after patch membrane rupture with a pipette containing WT h-tau (filled circles; Tau), endocytic rate was significantly prolonged (p < 0.05 compared to controls, open circles, repeated-measures two-way *ANOVA* with *post-hoc* Scheffe test, n = 5 from 5 slices), whereas exocytic magnitude remained similar to controls (p =0.45). 30 min after rupture, endocytic rate was further prolonged (p < 0.01) and exocytic magnitude became significantly less than controls (p < 0.05). Ca^2+^ current charge (Q_Ca_) remained unchanged throughout recording.

### Inhibition of SV endocytosis and synaptic transmission by WT h-tau requires *de novo* MT assembly

Since new MT assembly might take place after h-tau loading (Figure 1, Figure S1), we tested whether the tubulin polymerization blocker nocodazole might reverse the toxic effects of h-tau on SV endocytosis and synaptic transmission. In tubulin polymerization assays, nocodazole inhibited h-tau-dependent MT assembly in a concentration-dependent manner, with a maximal inhibition reached at 20 µM (Figure 3A). In presynaptic capacitance measurements, nocodazole (20 µM) co-loaded with h-tau (20 µM) fully prevented the h-tau toxicities on endocytosis (Figure 3B) and synaptic transmission (Figure 3C). Nocodazole alone (20 µM) had no effect on exo-endocytosis (Figure 3B) or EPSC amplitude (Figure 3C). It is highly likely that WT h-tau loaded in calyceal terminals newly assembled MTs, thereby impairing SV endocytosis and synaptic transmission.

**Figure 3.**
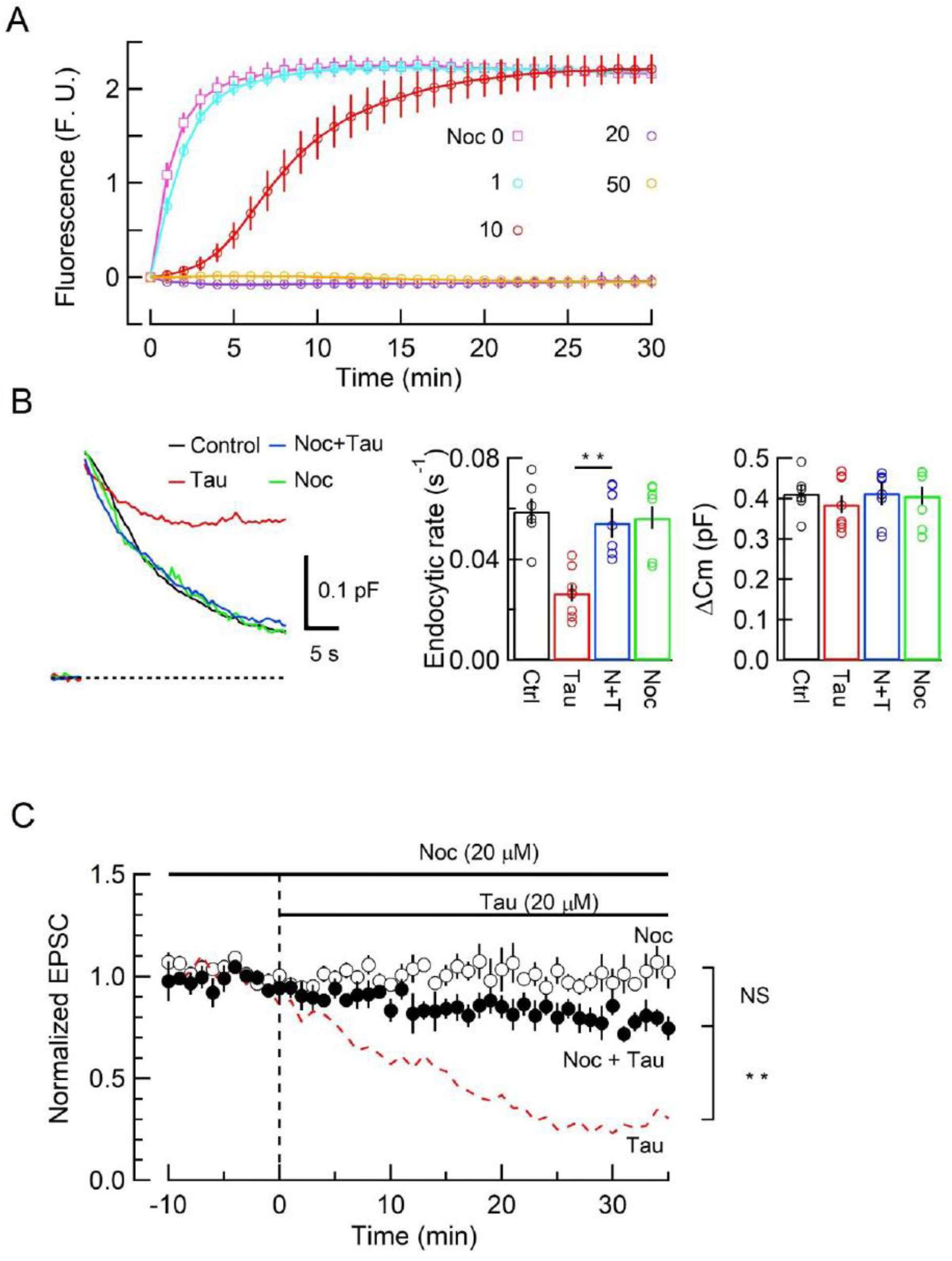
The MT assembly blocker nocodazole prevented tau-induced block of SV endocytosis and EPSC rundown. (A) Concentration-dependent inhibitory effects of nocodazole on MT assembly in tubulin polymerization assay. MT assembly by 0N4R h-tau (20 µM) in the absence (pink symbols and a fitting line) or presence of nocodazole at 1 µM (blue), 10 µM (red), 20 µM (purple) and 50 µM (orange). Data points and error bars in all graphs represent means and SEMs (n = 3). (B) Nocodazole prevented h-tau-induced block of SV endocytosis. Presynaptic membrane capacitance changes (superimposed traces) 25 min after loading h-tau alone (20 µM, red trace), h-tau and nocodazole (20 µM, blue), nocodazole alone (20 µM, green) and controls with no loading (black). Bar graphs indicate endocytic rates in non-loading controls (Ctr, black, 6 terminals from 6 slices), h-tau loaded terminals (Tau, red, 8 terminals from 8 slices), co-loading of nocodazole with h-tau (N+T, blue, 7 terminals from 7 slices) and nocodazole alone (Noc, green, 8 terminals from 8 slices). Nocodazole co-loading fully prevented endocytic block by h-tau (p < 0.01, between Tau and N-T) to control level (one-way ANOVA with Scheffe *post hoc* test). (C) Nocodazole prevented EPSC rundown caused by WT h-tau. Nocodazole (20 µM) co-loaded with WT h-tau (20 µM) prevented EPSC rundown (filled circles, 4 synapses from 4 slices, p < 0.01, unpaired t-test). Data of WT h-tau effect on EPSCs (Figure 1A) is shown as a red dashed line for comparison. Nocodazole alone (20 µM) had no effect on EPSC amplitude throughout (open circles, 4 synapses from 4 slices).

### WT h-tau assembles MTs and sequesters dynamins in calyceal terminals

The monomeric GTPases dynamin 1 and 3 play critical roles in the endocytic fission of SVs (50-52). Since dynamin is originally discovered as a MT-binding protein (39), we hypothesized that newly assembled MTs might trap free dynamins in cytosol. If this is the case, MT-bound form of dynamin would be increased. To test this hypothesis, we performed immunofluorescence microscopy and image analysis to quantify MTs and dynamin. After whole-cell infusion of h-tau into calyceal terminals, slices were chemically fixed and permeabilized to allow cytosolic free molecules such as tubulin monomers to be washed out of the terminal, thereby enhancing the signals from large structures such as MTs or bound molecules. Fluorescent h-tau antibody identified calyceal terminals loaded with WT h-tau (20 µM, Figure 4). Double staining with mouse β3-tubulin antibody revealed a 2.1-fold increase in MT signals in h-tau-loaded terminals, compared with those without h-tau loading (p = 0.01, n = 5, two-tailed unpaired t-test with Welch’s correction, Figure 4B). Triple labeling with dynamin antibodies further revealed a 2.6-fold increase in dynamin signal (p = 0.01, n = 5, two-tailed t-test with Welch’s correction, Figure 4B). These results suggest that soluble WT h-tau can assemble MTs in presynaptic terminals, thereby sequestering cytosolic dynamins that are indispensable for SV endocytosis.

**Figure 4.**
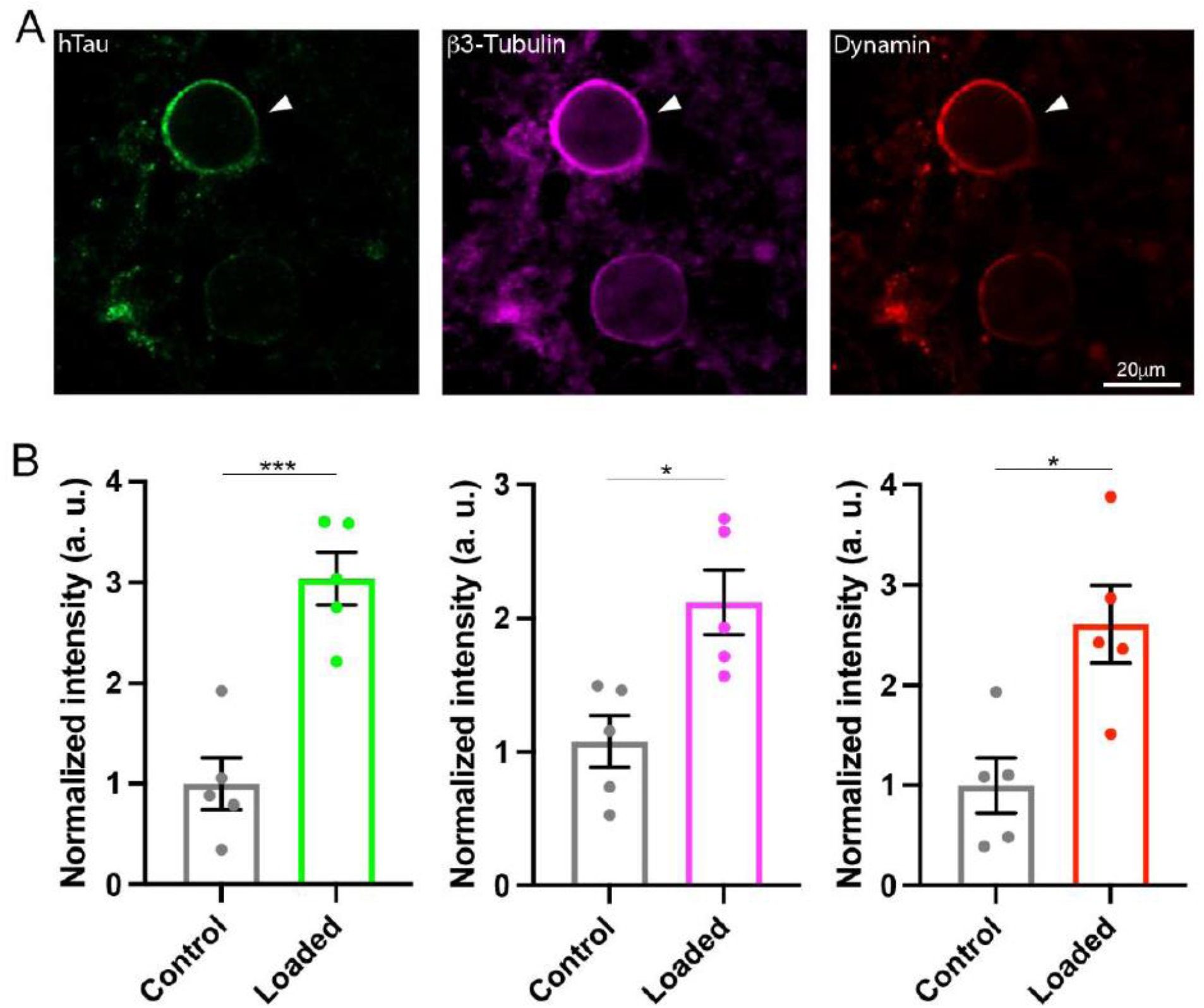
WT h-tau assembled MTs and increased bound-form dynamins in calyceal terminals. (A) Immunofluorescence images of brainstem slices showing loaded WT h-tau (green, left, arrowhead) labeled with anti h-tau/AlexaFluor-488 antibodies (green, left panel), newly assembled MTs labeled with anti β3-tubulin/AlexaFluor-647 antibodies (magenda, middle) and increased bound-form dynamin labeled with anti dynamin1/AlexaFluor-568 antibodies (red, right panel). (B) Bar graphs showing immunofluorescence intensities of h-tau (green), β3-tubulin (magenda) and dynamin (red) relative to controls with no loading (black bars). WT h-tau loading significantly increased β3-tubulin (p = 0.0105) and dynamin1 (p = 0.0109) intensity in terminals compared to control terminals without WT h-tau loading (n = 5 terminals from 5 slices for each data set, two-tailed unpaired t-test with Welch’s correction; * p < 0.05, *** p < 0.001).

### A microtubule-dynamin binding inhibitor peptide attenuates h-tau toxicities on SV endocytosis and synaptic transmission

To prevent toxic effects of h-tau on endocytosis and transmission, we searched for a dominant-negative (DN) peptide blocking MT-dynamin binding. Since the MT binding domain of dynamins is unknown, we synthesized 11 peptides from the pleckstrin-homology (PH) domain and 11 peptides from the proline-rich domain of dynamin 1 (Figure S2) and submitted them to the MT-dynamin1 binding assay. Out of 22 peptides, one peptide corresponding to the amino acid sequence 560-571 of PH domain, which we named “PHDP5”, significantly inhibited the MT-dynamin 1 interaction (Figure 5A). By SYPRO orange staining, dynamin 1 is found as a ∼100 kDa band, 1.7 ± 0.4 % in precipitates (ppts). In the presence of MT, dynamin 1 in ppts increased to 22.6 ± 2.4 %, indicating sequestration of dynamin 1 by MTs. When PHDP5 was added to MT and dynamin 1, dynamin1 in the ppt fraction decreased to 6.3 ± 2.4 %, indicating that PHDP5 works as a DN peptide for inhibiting MT-dynamin interactions (Figure 5A).

**Figure 5.**
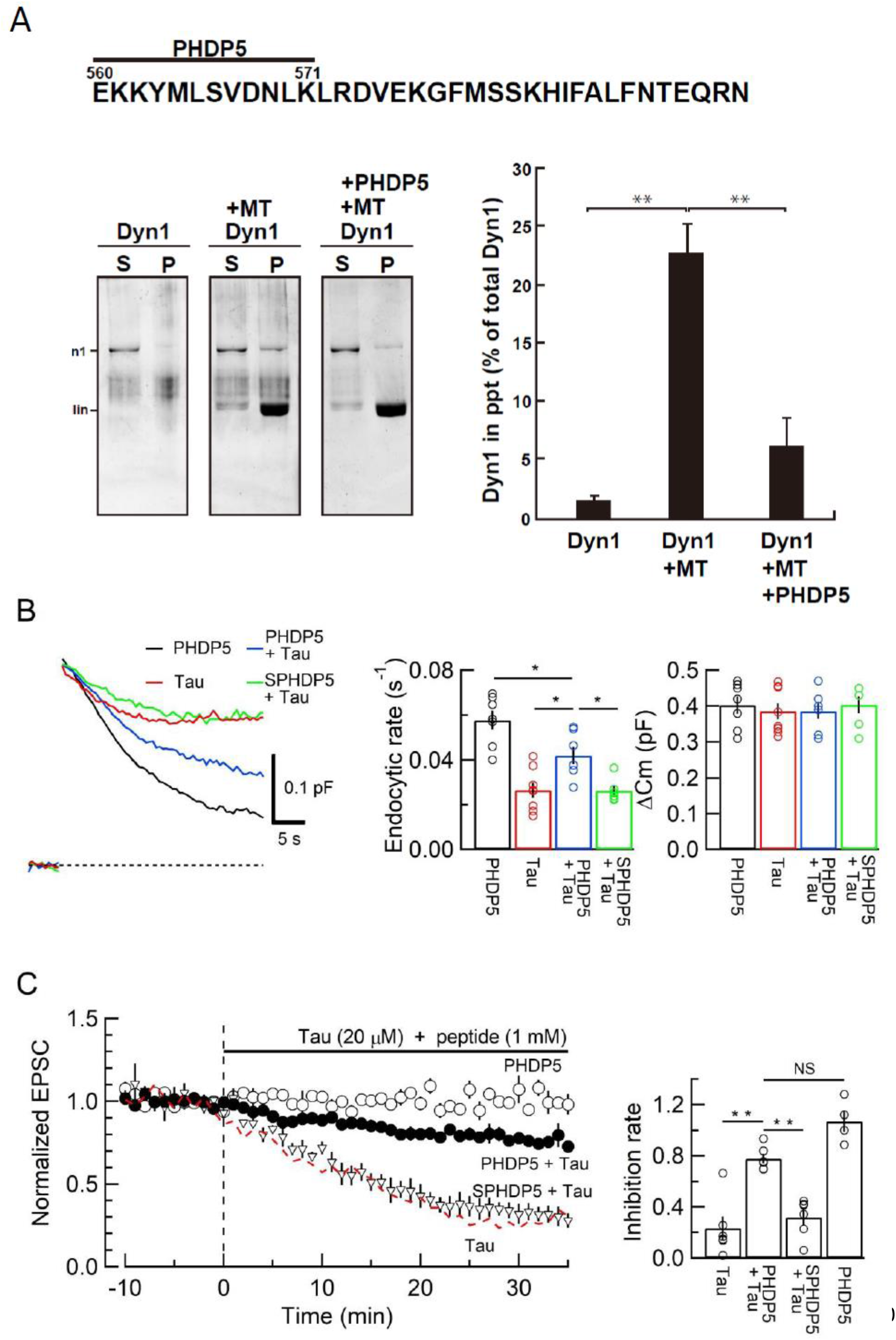
dynamin 1 PH domain peptide inhibited MT-dynamin 1 binding and prevented endocytic slowing and EPSC rundown caused by WT h-tau. (A) *Top*, Partial amino acid sequence of PH domain of mouse dynamin 1 indicating the sequence of the synthetic dodecapeptide PHDP5 (560-571). *Left*, SDS-PAGE of MT-dynamin 1 binding assay. S, supernatant; P, precipitates. Dyn1, dynamin 1. *Right*, quantification of MT-dynamin 1 interaction. The bars indicate the percentage of dynamin 1 found in precipitates relative to total amount. PHDP5 significantly inhibited MT-dynamin 1 interaction (**<0.01, n = 3). (B) Presynaptic membrane capacitance records (superimposed) after loading h-tau alone (20 µM, red trace, taken from Figure 4B), h-tau co-loaded with DPHP5 (0.25 mM, blue) or scrambled DPHP5 (SDPHP5, green). DPHP5 alone (0.25 mM, black trace, 7 terminals from 7 slices) had no effect on capacitance changes compared to non-loading terminal controls (taken from Figure 3B). Bar graphs of endocytic rates (middle panel) indicate significant difference (p < 0.05, 7 terminals from 7 slices) between tau (red bar, 8 terminals from 8 slices) and DPHP5+tau (blue, 7 terminals from 7 slices) as well as between SDPHP5 + tau (blue) and DPHP5+ tau (green, n = 6 terminals from 6 slices). The magnitudes of exocytic capacitance changes were not significantly different between the groups, recorded 25 min after rupture). (C) DPHP5 attenuated h-tau induced EPSC rundown. The EPSC rundown after h-tau infusion (20 µM, red dashed line; data taken from Fig 1A) was attenuated by DPHP5 (1 mM) co-loaded with h-tau (filled circles) but not by scrambled DPHP5 peptide (SDPHP5, open triangles, 1 mM). DPHP5 alone (1 mM, open circles) had no effect on EPSC amplitude throughout. Bar graphs indicate EPSC amplitude (normalized to that before infusion) 30 min after infusion. Significant difference (p < 0.01) between tau and tau + DPHP5, between tau + DPHP5 and tau + SDPHP5. The difference between DPHP5 alone and DPHP5 + tau was not significant (p = 0.09), indicating the partial antagonistic effect of DPHP5 against h-tau-induced EPSC rundown.

Loading of PHDP5 (0.25 mM) alone in calyceal terminals had no effect on exo-endocytic capacitance changes, but when co-loaded with WT h-tau (20 µM), it significantly attenuated the h-tau-induced endocytic slowing (p < 0.05, Figure 5B). Scrambled PHDP5 peptide (0.25 mM) loaded as a control had no effect on h-tau-induced endocytic slowing. Similar to its effect on capacitance changes, intra-terminal infusion of PHDP5 alone (1 mM) did not affect EPSC amplitude, but when co-loaded with WT h-tau (20 µM), significantly attenuated the inhibitory effect of h-tau on EPSC amplitude (p < 0.01, Figure 5C). Co-infusion of scrambled PHDP5 (1 mM) with h-tau (20 µM) did not affect the h-tau-induced EPSC rundown (p = 0.46). These results further support that WT h-tau causes dynamin deficiency via new assembly of MTs thereby impairing SV endocytosis and synaptic transmission. These results also highlight PHDP5 as a potential therapeutic tool for rescuing synaptic dysfunctions associated with AD or PD.

## Discussion

Using the calyx of Held in brainstem slices as an AD model for dissecting mammalian central excitatory synaptic transmission, we demonstrated that intra-terminal loading of WT h-tau impairs vesicle endocytosis and synaptic transmission via *de novo* MT assembly. Previous over-expression studies in cultured cells reported MT assembly by WT tau (36, 53, 54) or phosphorylated tau (34, 55). Compared with overexpression, our whole-cell method allows targeted loading of molecules in presynaptic terminals at defined concentrations because of a large pipette-to-cell volume ratio (44). In postmortem brain tissue homogenates from AD patients, soluble tau content is estimated as 6 ng/μg of protein, which is 8 times higher than controls (56). Assuming protein contents in brain homogenate as 10%, 60 kDa tau concentration in AD patients’ brain is estimated as 10 µM. Since elevation of soluble tau concentration likely occurs mainly in axons and axon terminal compartments of neurons, soluble tau concentration in AD patients in presynaptic terminals can be higher than that. Our results at the calyx of Held suggest that excitatory synaptic transmission, in general, can be significantly impaired in such situations. In fact, the magnitude of EPSC rundown after WT h-tau loading is comparable to that caused by the clinical dose of general anesthetic isoflurane at the calyx of Held in slice (48). In AD, tau pathology starts from the locus coeruleus in the brainstem and undergoes trans-synaptic propagation to hippocampal and neocortical neurons (57). Present results in our model synapse suggest that synaptic functions in such tau-propagation pathways can be severely affected at the early stage of AD.

Membrane capacitance measurements at the calyx of Held revealed the primary target of WT h-tau toxicity as SV endocytosis. Endocytic slowing impairs SV recycling and reuse, thereby inhibiting SV exocytosis, particularly in response to high-frequency stimulations (49). The toxic effects of h-tau on SV endocytosis and synaptic transmission were prevented by nocodazole co-application. Together with the lack of toxicity of del-MTBD and toxic effects of taxol on synaptic transmission, these results suggest pathological roles of over-assembled MTs. Like WT h-tau, intra-terminal loading of WT α-synuclein slows SV endocytosis and impairs fidelity of high-frequency neurotransmission at the calyx of Held (46). α-Synuclein toxicities can be rescued by blocking MT assembly with nocodazole or a photosensitive colchicine derivative PST-1. Thus, a common mechanism likely underlies synaptic dysfunctions in AD and PD. Compared with α-synuclein, h-tau toxicity is much stronger on endocytosis as well as on synaptic transmission. Thus, abnormal elevation of endogenous molecules beyond homeostatic level may cause AD and PD symptoms, like many other human diseases.

Although the GTPase dynamin is a well-known player in endocytic fission of SVs (50, 52), it was originally discovered as a MT-binding protein (39). Subsequent studies indicated that this interaction upregulates dynamin’s GTPase activity (58, 59) and can induce MT instability with dynamin 2 (60) or stabilizes MT bundle formation with dynamin 1 (61). However, the binding domain of dynamin remained unidentified. In this study, calyceal terminals loaded with WT h-tau showed a prominent increase in immunofluorescence signal intensity corresponding to bound dynamins. This was associated with an elevation in intra-terminal MTs, suggesting that newly assembled MTs induced by loaded h-tau sequestered cytosolic dynamins. These results well explain impairments of SV endocytosis by intra-terminal h-tau loading. Through synthetic peptide screening, we found that a dodecapeptide from dynamin 1 PH domain significantly inhibited the MT-dynamin interaction. This peptide PHDP5, which is ∼80 % homologous to dynamin 3, another isoform involved in vesicle endocytosis (51), rescued endocytic impairments and EPSC rundown induced by intra-terminal WT h-tau. Hence, MTs over-assembled by soluble WT h-tau proteins likely sequester free dynamins in presynaptic terminals, thereby blocking SV endocytosis and synaptic transmission, at least in this slice model. This dynamin sequestration mechanism by newly assembled MTs may also underlie the toxic effect of α-synuclein on SV endocytosis (46) in PD.

Unlike WT h-tau, the FTDP-linked mutant tau does not affect SV endocytosis (32), but binds to both actin filaments (62) and the SV transmembrane protein synaptogyrin (31), thereby immobilizing SVs (31, 32). WT-tau can also bind to synaptogyrin (31), but cannot bind to F-actins because of a difference in the MT-binding regions between FTDP mutant and WT tau (53, 63). However, WT h-tau can bind to various other macromolecules and organelles such as MTs, neurofilaments and ribosomes (64) as well as to synaptogyrin, thereby possibly immobilizing SVs. Recycling transport of SVs impaired by this mechanism might additionally contribute to the rundown of synaptic transmission remaining unblocked by the MT-dynamin blocker peptide. In the absence of a powerful tool for alleviating symptoms associated with AD or PD, the calyx of Held slice model might provide a platform upon which therapeutic tools for rescuing synaptic dysfunctions can be pursued. The combination of this slice model with animal models could provide a new pathway toward rescuing neurological disorders.

## Materials and Methods

### Animals

All experiments were performed in accordance with the guidelines of the Physiological Society of Japan and animal experiment regulations at Okinawa Institute of Science and Technology Graduate University.

### Recombinant human tau preparation

Human tau (h-tau) lacking the MT-binding domain (amino acid 244 to 367, del-MTBD) were produced by site-directed mutagenesis as previously reported (40). Wild-type (WT) and del-MTBD mutant h-tau of 0N4R isoform were expressed in *E. coli*. (BL21/DE3) and purified as described previously (65) with minor modifications. Briefly, harvested bacteria expressing recombinant tau were lysed in homogenization buffer (50 mM PIPES, 1 mM EGTA, 1 mM DTT, 0.5 mM PMSF, and 5 µg/ml Leupeptin, pH6.4), sonicated and centrifuged at 27,000 xg for 15 min. Supernatants were charged onto phosphocellulose column (P11, Whatman). After washing with homogenization buffer containing 0.1 M NaCl, h-tau-containing fractions were eluted by the buffer containing 0.3 M NaCl. Subsequently, the proteins were precipitated by 50 % saturated ammonium sulfate and re-solubilized in homogenization buffer containing 0.5 M NaCl and 1% 2-mercaptoethanol. After incubation at 100 °C for 5 min, heat stable (soluble) fractions were obtained by centrifugation at 21,900 xg, and fractionated by reverse phase high-performance liquid chromatography (RP-HPLC) using Cosmosyl Protein-R (Nacalai tesque Inc.). Aliquots of h-tau containing fractions were lyophilized and stored at -80 °C. Purified h-tau proteins were quantified by SDS-PAGE followed by Coomassie Brilliant Blue staining.

### Purification of recombinant human dynamin1 protein

His-tagged human dynamin 1 was expressed using the Bac-to-Bac baculovirus expression system (Thermo Fisher Scientific, Waltham, MA, USA) and purified as described previously (66). The purified dynamin solutions were concentrated using Centriplus YM50 (cat#4310; Merck-Millipore, Darmstadt, Germany).

### Microtubule polymerization assay

Effects of tau and nocodazole on MT polymerization were tested using a Tubulin Polymerization Assay (Cytoskeleton Inc., Denver, CO). Briefly, purified WT or del-MTBD mutant h-tau (10 µM) were mixed with porcine tubulin (20 µM) in an assembly buffer at 37 °C. Nocodazole was added to the mixture at 0 min of incubation. MT polymerization was fluorometrically assayed (excitation at 360 nm, emission at 465 nm) using Infinit F-200 Microplate Reader (TECAN, Männedorf / Switzerland) at 1 min intervals for 30 min. After incubation, resultant solutions were subjected to centrifugation at 100,000 xg for 15 min at 20°C. Supernatants (free tubulin fraction) and pellets (microtubule fraction) were subjected to SDS-PAGE to quantify the amount of tubulin assembled into MTs.

### Peptide synthesis and LC-MS/MS analysis

The peptides were synthesized through conventional 9-fluorenylmethyloxycarbonyl (Fmoc) solid-phase peptide synthesis (SPPS), onto preloaded Fmoc-alanine TCP-resins (Intavis Bioanalytical Instruments) using automated peptide synthesizer ResPep SL (Intavis Bioanalytical Instruments). All Fmoc-amino acids were purchased from Watanabe Chemical Industries and prepared at 0.5 M in N-methyl pyrrolidone (NMP, Wako Pure Chemical Industries). After synthesis, peptides were cleaved with (v/v/v) 92.5% TFA, 5% TIPS and 2.5% water for 2 h, precipitated using t-butyl-methyl-ether at -30° C, pelleted and resuspended in water before lyophilization (EYELA FDS-1000) overnight. All synthesized peptides’ purity and sequence were then confirmed by LC-MS/MS using a Q-Exactive Plus Orbitrap hybrid mass spectrometer (Thermo Scientific) equipped with Ultimate 3000 nano-HPLC system (Dionex), HTC-PAL autosampler (CTC Analytics), and nanoelectrospray ion source.

### MT-dynamin binding assay

Microtubule binding assay was performed using Microtubule Binding Protein Spin Down Assay Kit (cat#BK029, Cytoskeletion Inc., Denver, CO, USA). Briefly, 20 μl of 5 mg/ml tubulin in general tubulin buffer (GTB; 80 mM PIPES pH7.0, 2 mM MgCl_2_, 0.5 mM EGTA) supplemented with 1 mM GTP were polymerized by adding 2 μl of cushion buffer (80 mM PIPES pH7.0, 1 mM MgCl_2_, 1 mM EGTA, 60% glycerol), and incubated at 35°C for 20 min. Microtubules were stabilized with 20 μM Taxol. Taxol-stabilized microtubules (2.5 μM) and dynamin 1 (1 μM) were incubated in GTB with or without 1 mM peptide at room temperature for 30 min. After incubation, the 50 μl of mixture was loaded on top of 100 μl cushion buffer supplemented with 20 μM Taxol, and then centrifuged at 100,000 xg for 40 min at room temperature. After the ultracentrifugation, 50 μl of supernatant was taken and mixed with 10 μl of 5×sample buffer. The resultant pellet was resuspended with 50 μl of 1×sample buffer. Twenty μl of each sample was analyzed by SDS-PAGE and stained with SYPRO Orange. Protein bands were visualized using FLA-3000 (FUJIFILM Co.LTD, Tokyo, Japan).

### Immunocytochemical analysis

The following primary antibodies were used: anti-β3-tubulin (Synaptic System, #302304), anti-human Tau (BioLegend, #806501), anti-dynamin (Invitrogen, PA1-660). Secondary antibodies were goat IgG conjugated with Alexa Fluor 488, 568, or 647 (Thermo Fisher Scientific). Acute brainstem slices (175 μm in thickness) were fixed with 4% paraformaldehyde in PBS for 30 min at 37 °C and overnight at 4 °C. On the following day, slices were rinsed three times in PBS, permeabilized in PBS containing 0.5% Triton X-100 (Tx-100; Nacalai Tesque) for 30 min and blocked in PBS containing 3% bovine serum albumin (BSA; Sigma-Aldrich) and 0.05% Tx-100 for 45 min. Slices were incubated overnight at 4 °C with primary antibody diluted in PBS 0.05% Tx-100, 0.3% BSA. On the next day, slices were rinsed three times with PBS containing 0.05% Tx-100 for 10 min and incubated with corresponding secondary antibody diluted in PBS 0.05% Tx-100, 0.3% BSA for 1 h at room temperature (RT). Slices were further rinsed three times in PBS 0.05% Tx-100 for 10 min and finally washed in PBS for another 10 min. Finally, slices were mounted on glass slides (Matsunami) using liquid mounting medium (Ibidi) and sealed using nail polish. Confocal images were acquired on laser scanning microscopes (LSM780 or LSM900, Carl Zeiss) equipped with a Plan-apochromat 63x oil-immersion objective (1.4 NA) and 488, 561, and 633 nm excitation laser lines. For quantifying fluorescence intensity levels, the region of interest was delimited around calyceal terminals, and background fluorescence was subtracted using ImageJ software.

### Slice Electrophysiology

After killing C57BL6N mice of either sex (postnatal day 13-15) by decapitation under isoflurane anesthesia, brainstems were isolated and transverse slices (175 μm thick) containing the medial nucleus of the trapezoid body (MNTB) were cut using a vibratome (VT1200S, Leica) in ice-cold artificial cerebrospinal fluid (aCSF, see below) with reduced Ca^2+^ (0.1 mM) and increased Mg^2+^ (3 mM) concentrations or sucrose-base aCSF (NaCl was replaced to 300 mM sucrose, concentrations of CaCl_2_ and MgCl_2_ was 0.1 mM and 6 mM, respectively). Slices were incubated for 1h at 36-37 °C in standard aCSF containing (in mM); 125 NaCl, 2.5 KCl, 26 NaHCO_3_, 1.25 NaHPO_4_, 2 CaCl_2_, 1 MgCl_2_, 10 glucose, 3 myo-inositol, 2 sodium pyruvate, and 0.5 sodium ascorbate (pH 7.4 when bubbled with 95 % O_2_ and 5 % CO_2_, 310-320 mOsm), and maintained thereafter at room temperature (RT, 24-28 °C).

Whole-cell recordings were made using a patch-clamp amplifier (Multiclamp 700A, Molecular Devices, USA for pair recordings and EPC-10 USB, HEKA Elektronik, Germany for presynaptic capacitance measurements) from the calyx of Held presynaptic terminals and postsynaptic MNTB principal neurons visually identified with a 60X or 40X water immersion objective (LUMPlanFL, Olympus) attached to an upright microscope (Axioskop2, Carl Zeiss, or BX51WI, Olympus, Japan). Data were acquired at a sampling rate of 50 kHz using pClamp (for Multiclamp 700A) or Patchmaster software (for EPC-10 USB) after online filtering at 5 kHz. The presynaptic pipette was pulled for the resistance of 7-10 MΩ and had a series resistance of 14-20 MΩ, which was compensated by 70 % for its final value to be 7 MΩ. Resistance of the postsynaptic pipette was 5-7 MΩ, and its series resistance was 10-25 MΩ, which was compensated by up to 75 % to a final value of 7 MΩ. The aCSF routinely contained picrotoxin (10 μM) and strychnine hydrochloride (0.5 μM) to block GABA_A_ receptors and glycine receptors, respectively. Postsynaptic pipette solution contained (in mM): 130 CsCl, 5 EGTA, 1 MgCl_2_, 5 QX314-Cl, 10 HEPES (adjusted to pH 7.3–7.4 with CsOH). The presynaptic pipette solution contained (in mM); 105 K methanesulfonate, 30 KCl, 40 HEPES, 0.5 EGTA, 1 MgCl_2_, 12 phosphocreatine (Na salt), 3 ATP (Mg salt), 0.3 GTP (Na salt) (pH 7.3-7.4 adjusted with KOH, 315-320 mOsm). WT h-tau protein or del-MTBD tau mutant was loaded using the pipette perfusion technique (42, 67).

Membrane capacitance (C_m_) measurements were made from calyx of Held presynaptic terminals in the whole-cell configuration at RT (47, 49). Calyceal terminals were voltage-clamped at a holding potential of -80 mV, and a sinusoidal voltage command (1 kHz, 60 mV in peak-to-peak amplitude) was applied. To isolate presynaptic voltage-gated Ca^2+^ currents (I_Ca_), the aCSF contained 10 mM tetraethylammonium chloride, 0.5 mM 4-aminopyridine, 1 µM tetrodotoxin, 10 µM bicuculline methiodide and 0.5 µM strychnine hydrochloride. The presynaptic pipette solution contained (in mM): 125 Cs methanesulfonate, 30 CsCl, 10 HEPES, 0.5 EGTA, 12 disodium phosphocreatine, 3 MgATP, 1 MgCl_2_, 0.3 Na_2_GTP (pH 7.3 adjusted with CsOH, 315-320 mOsm). Tau solution was backfilled into the pipette after loading the tau-free pipette solution. Care was taken to maintain series resistance < 16 MΩ to allow dialysis of the terminal with pipette solution. Recording pipette tips were coated with dental wax to minimize stray capacitance (4-5 pF). Single square pulse (−80 to 10 mV, 20 ms duration) was used to induce presynaptic I_Ca_. Membrane capacitance changes within 450 ms of square-pulse stimulation were excluded from analysis to avoid contamination by conductance-dependent capacitance artifacts (49). To avoid the influence of capacitance drift on baseline, we removed data when the baseline drift measured 0-10 s before stimulation was over 5 fFs^-1^. When the drift was 1-5 fFs^-1^, we subtracted a linear regression line of the baseline from the data for the baseline correction. The endocytic rate was calculated from the slope of the normalized C_m_ changes during the initial 10 s after the stimulation.

### Data analysis and statistics

Data were analyzed using IGOR Pro 6 (WeveMatrics), Excel 2016 (Microsoft), and StatPlus (AnalystSoft Inc) and KaleidaGraph for Macintosh, version 4.1 (Synergy Software Inc., Essex Junction, VT, USA). All values are given as mean ± S.E.M. Differences were considered statistically significant at *p* < 0.05 in paired or unpaired *t*-tests, one-way *ANOVA* with Scheffe *post-hoc* test and repeated-measures two-way *ANOVA* with *post-hoc* Scheffe test.

## Acknowledgments

We thank Yasuo Ihara, Nobuyuki Nukina and Takeshi Sakaba for comments and Patrick Stoney for editing this paper. We also thank Shota Okuda and Mikako Matsubara for their contributions in the early stage of this study, and Satoko Wada-Kakuda for technical assistant with *in vitro* analysis of tau. This research was supported by funding from Okinawa Institute of Science and Technology and from Technology (OIST) and Core Research for the Evolutional Science and Technology of Japan Science and Technology Agency (CREST) to T.T., and by Scientific Research on Innovative Areas to T.M (Brain Protein Aging and Dementia Control 26117004).

## Supplementary Information for

**Other supplementary materials for this manuscript include the following:**

N/A

**Fig. S1.**
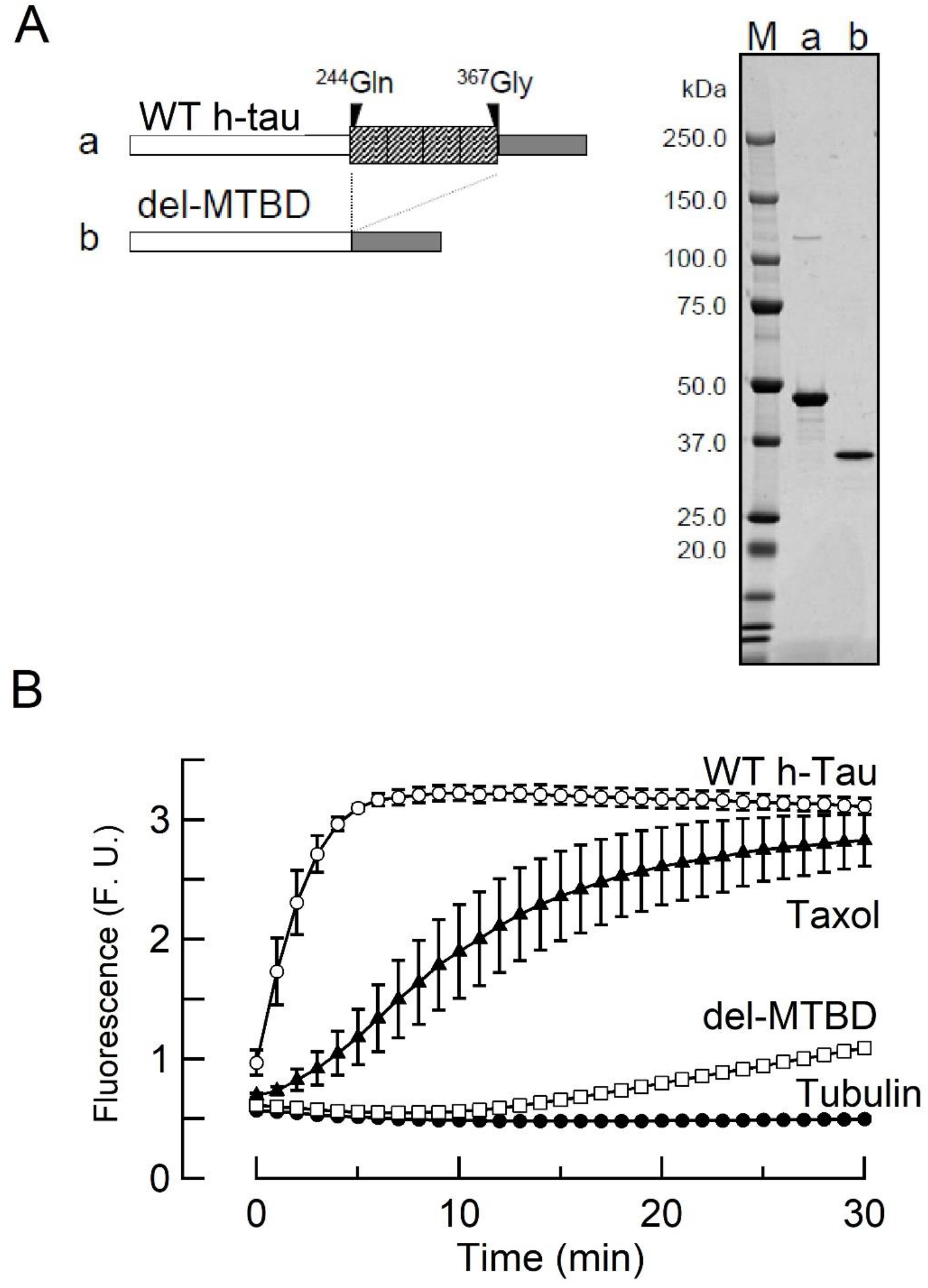
Tubulin polymerization assay for purified ON4R WT h-tau and MT-binding region-deleted mutant. (A) *Left panel*, schematic drawings of WT 0N4R h-tau (a) and h-tau deletion mutant lacking the MT-binding region (del-MTBD, b). *Right panel*, purified recombinant WT 0N4R h-tau (a) and del-MTBD (b) in SDS-PAGE with molecular markers (M) on the left lane. (B) *In vitro* tubulin polymerization assay, showing MT assembly by WT h-tau (10 µM, open circles) or taxol (1 µM, filled triangles), but not by del-MTBD (10 µM, open squares) or tubulin alone (filled circles). Data points and bars represent means and SEMs (n =3).

**Fig. S2.**
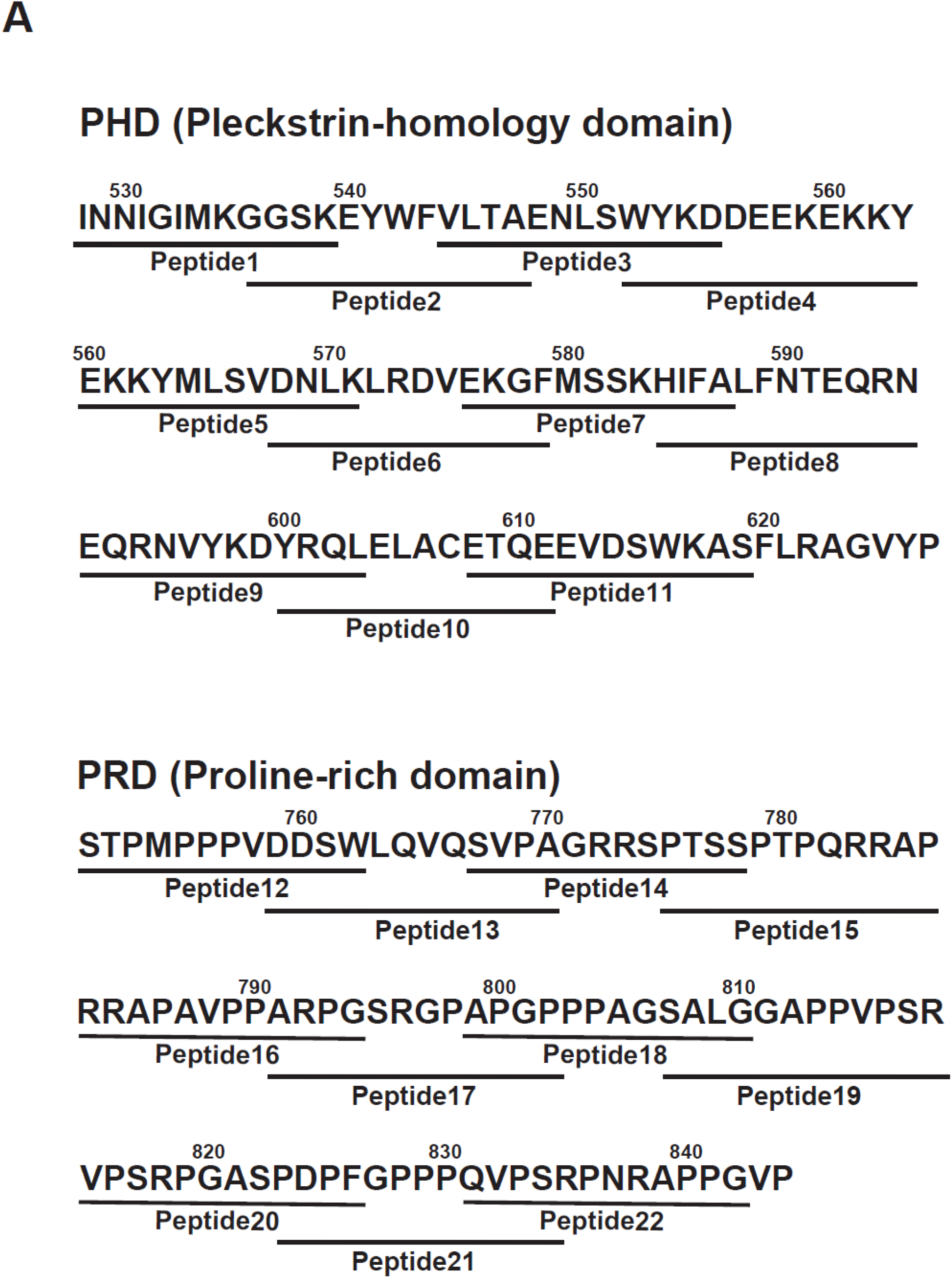
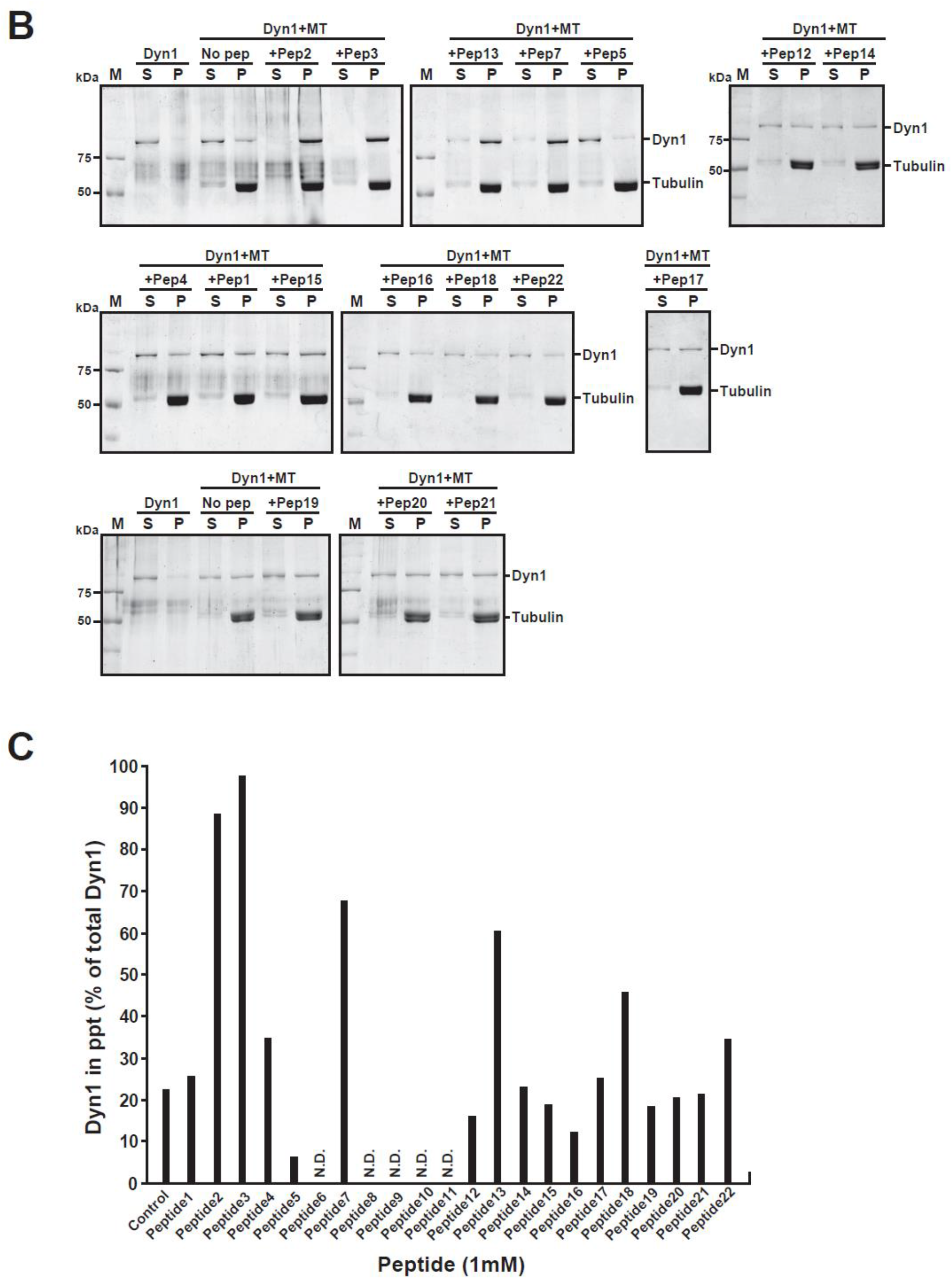
PHDP5 strongly inhibited dynamin 1 binding to microtubules. (A) Peptide sequences in mouse dynamin 1 used in the DN peptide screening. In total, 24 peptides covering the PH domain and the proline-rich domain were synthesized. (B) SDS-PAGE of the microtubule binding assay, showing that PHDP5 strongly inhibits the MT-dynamin 1 binding (“+Pep5” in the top right panel). Since 5 peptides (peptide6, peptide8, peptide9, peptide10, peptide11) were insoluble in water or dimethyl sulfoxide, their effects were not tested. (C) Effects of the synthetic peptides at 1 mM on MT-dynamin1 binding. Bar graphs indicate the percentage of dynamin 1 found in precipitates relative to total amount.

